# The developing mouse coronal suture at single-cell resolution

**DOI:** 10.1101/2021.02.24.432636

**Authors:** D’Juan T. Farmer, Hana Mlcochova, Yan Zhou, Nils Koelling, Guanlin Wang, Neil Ashley, Robert E Maxson, Andrew O. M. Wilkie, J Gage Crump, Stephen R.F. Twigg

## Abstract

Sutures separate the flat bones of the skull and enable coordinated growth of the brain and overlying cranium. To uncover the cellular diversity within sutures, we generated single-cell transcriptomes and performed extensive expression validation of the embryonic murine coronal suture. We identify *Erg* and *Pthlh* as markers of osteogenic progenitors in sutures, and distinct pre-osteoblast signatures between the bone fronts and periosteum. In the ectocranial layers above the suture, we observe a ligament-like population spanning the frontal and parietal bones. In the dura mater underlying the suture, we detect a chondrocyte-like signature potentially linked to cartilage formation under pathological conditions. Genes mutated in coronal synostosis are preferentially expressed in proliferative osteogenic cells, as well as meningeal layers, suggesting discrete cell types that may be altered in different syndromes. This single-cell atlas provides a resource for understanding development of the coronal suture, the suture most commonly fused in monogenic craniosynostosis.

## Introduction

Cranial sutures are fibrous joints between calvarial bones that act as zones of bone growth and absorbers of physical forces^1^. They comprise the leading edges of abutting calvarial bones separated by mesenchymal tissue. New bone forms by intramembranous ossification in response to expansion of the underlying brain^1,2,3^. Growth of the bony skull requires the proliferation and osteogenic differentiation of progenitor cells, as well as the maintenance of sufficient undifferentiated cells in the suture to ensure continued bone growth during fetal and postnatal stages. Environmental and/or genetic insults that disrupt the delicate balance of proliferation and differentiation result in premature fusion of cranial sutures, a condition known as craniosynostosis^4,5^.

The coronal suture, which separates the frontal and parietal bones, is the suture most commonly affected in monogenic craniosynostosis^6^. The mouse has proven an effective model for the study of coronal synostosis^7,8,9,10^. During cranial development, the coronal suture and closely associated tissues are derived from three distinct populations: the supraorbital mesenchyme which will give rise to the calvarial bones and suture mesenchyme, the meningeal mesenchyme, and non-osteogenic early migrating mesenchyme^11^. The meninges form between the brain and calvaria and are essential for development of both^12^. Coronal suture mesenchyme, derived largely from the mesoderm, forms a boundary between the neural crest-derived frontal and mesoderm-derived parietal bones^13,14^. Embryonic suture mesenchyme originates from *Gli1*-expressing cells that migrate away from the paraxial cephalic mesoderm at embryonic day (E) 7.5^15^ and expand apically to sit between the lateral dermal mesenchyme and medial meningeal layers from E12.5 onwards^11^. Whereas these macroscopic developmental steps are well established, the cellular composition of the developing coronal suture remains poorly understood. Markers that label skeletal stem cells within postnatal sutures have been identified^16,17,18,19^, yet none of these markers identify a distinct skeletal progenitor cell population at embryonic stages. Given recent reports that embryonic progenitor dysfunction precedes craniosynostosis^20^, identifying cell type diversity in early forming sutures will be critical for understanding the etiology of this birth defect. A better understanding of embryonic osteogenic and non-osteogenic populations will also inform how the meninges and ectocranial layers contribute to suture patency^7,21,22^.

To build a cell atlas of the embryonic coronal suture, we combined single-cell transcriptomics with highly resolved in situ analysis to catalogue the cell types present in murine coronal sutures at E15.5 and E17.5. In the ectocranial compartment, we uncovered multiple layers of distinct cell types, including a ligament-like population connecting the lateral aspects of the frontal and parietal bones. Within the multiple layers of the meninges, we revealed an outer dura mater population with a chondrogenic signature, suggesting a latent capacity for chondrocyte differentiation. In the osteogenic population, pseudotime analysis revealed a putative *Erg*^+^/*Pthlh*^+^ progenitor that we found to be located in the suture and along the leading edges of the bones. These progenitors fed into two distinct pre-osteoblast trajectories, one concentrated at the growing bone tips and the other localized along the periosteum more distant from the suture. Comparative expression analysis between genes associated with coronal or midline synostosis highlighted the selective expression of many coronal synostosis genes, including *Twist1* and *Tcf12*, within proliferative osteogenic cells, but also within ectocranial and meningeal layers, suggesting heterogenous etiologies for coronal synostosis. We also detected potential ligand-receptor interactions of neighboring ectocranial and meningeal layers with osteogenic cells within the suture, in particular the proliferative osteogenic population. This is in agreement with previous studies showing roles for the ectocranial mesenchyme and meninges in regulating suture patency^11,12^. This single-cell atlas reveals diversity within the developing coronal suture, some of the earliest potential markers for suture-resident osteogenic progenitors, and potential interactions between osteogenic and non-osteogenic populations likely important for proper skull expansion.

## Results

### Diverse mesenchymal heterogeneity captured by single-cell sequencing

To understand the cellular composition of the embryonic coronal suture, we performed single-cell RNA sequencing at E15.5 and E17.5 on dissected coronal sutures, including small amounts of frontal and parietal bone, after removing the skin and brain (Fig. 1a). We filtered using Seurat 3 R-Package^23^ and obtained 8279 cells at E15.5 (median of 2460 genes per cell) and 8682 cells at E17.5 (median of 3200 genes per cell) (Fig. 1b). We identified 14 cell clusters through unsupervised graph clustering of the two datasets combined (Supplementary Table 1). Osteogenic and mesenchymal cell types were identified based on the expression of broad mesenchyme/fibroblast (*Col1a1*) and osteoblast (*Sp7*) markers (Fig. 1b, dotted line; Supplementary Fig. 1a-b). The identities of clusters outside the osteogenic/mesenchymal subset were resolved using previously reported markers, and included chondrocytes, myeloid cells, mast cells, lymphocytes, pericytes, osteoclasts, endothelial cells, neurons, and glia (Fig. 1b, c). All the major cell types in our analysis were present at E15.5 and E17.5 (Supplementary Fig. 2a). Chondrocytes were especially abundant at E15.5 (Supplementary Fig. 2b), highlighting the close proximity of the E15.5 coronal suture to the chondrocranium. Myeloid cells were more abundant at E17.5, consistent with reports of increased myeloid differentiation during late embryonic stages^24^ (Supplementary Fig. 2b).

**Figure 1.**
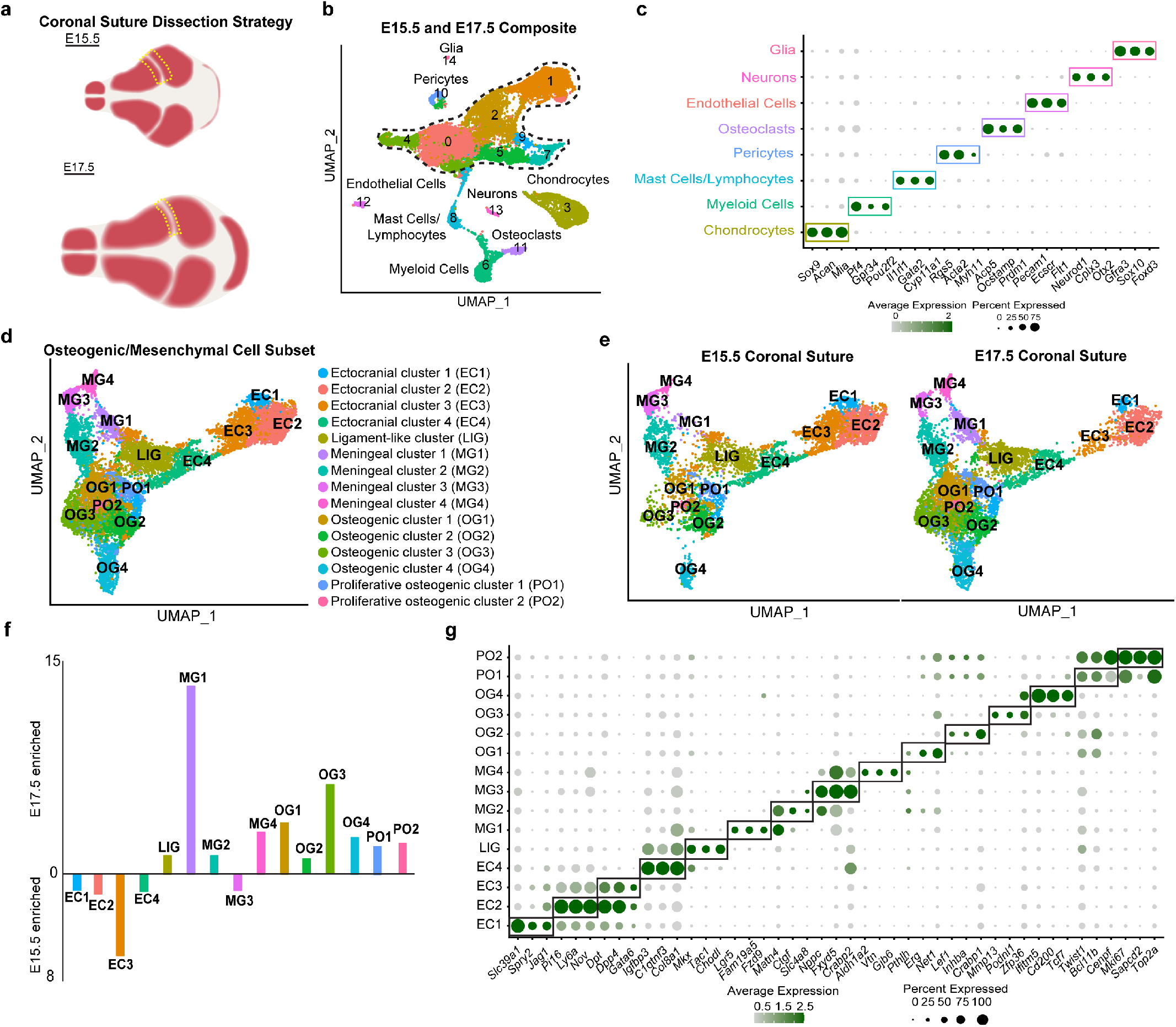
Single-cell RNA-sequencing analysis of the E15.5 and E17.5 coronal suture. **a** Schematic of dissection strategy for E15.5 and E17.5 coronal sutures. Yellow dashed lines outline dissected regions. **b** Uniform Manifold Approximation and Projection (UMAP) plot of integrated E15.5 (8279 cells) and E17.5 (8682 cells) datasets. Osteogenic and mesenchymal cell types outlined by dashed lines. **c** Dot plot depicting selected markers (determined by adjusted p-value) enriched for each ancillary cell type outside of the osteogenic and mesenchymal population. **d** UMAP analysis of re-clustered osteogenic and mesenchymal subset outlined in (b) resolves 14 clusters. **e** The osteogenic and mesenchymal subset separated by developmental stage. **f** Graphical depiction of the cluster proportions from the osteogenic/mesenchymal subset, plotted as ratio between E15.5 and E17.5 cells within each cluster. **g** Dot plot showing markers enriched for each cluster within the osteogenic/mesenchymal subset.

To analyse the osteogenic and mesenchymal cell types that might comprise and support the coronal suture, we re-clustered the osteogenic/mesenchymal population and obtained 14 clusters present at both E15.5 and E17.5 (Fig. 1d-f). Analysis of enriched genes for each cluster allowed us to assign probable identities to each cell type (Fig. 1g, Supplementary Table 2), which we validated by in situ hybridization as described below. We observed one ectocranial cluster strongly over-represented at E15.5 and a meningeal cluster over-represented at E17.5. Osteogenic cells were also more abundant in the E17.5 dataset, although it is unclear whether this reflects true biological differences versus differing cell capture between the dissections (Fig. 1a, f).

### Diversity of meningeal layers below the coronal suture

The meninges are involved in the development of the calvaria and underlying brain that they separate^12^. They comprise dura mater, arachnoid mater, and pia mater. To determine the identity of meningeal tissues included in our dissections, we utilized a recent transcriptomic study of murine E14 meninges as a guide^25^. Markers associated with the pia mater (*Ngfr*, *Lama1*, *Rdh10*) were not co-enriched in any of our clusters, consistent with the pia mater being removed with the brain during dissections (Fig. 2a). In contrast, markers enriched within the arachnoid mater (*Aldh1a2*, *Cldn11*, and *Tbx18*) were abundant in MG4, and dura mater markers (*Gja1*, *Fxyd5*, and *Crabp2*) in MG3 and MG4, and to a lesser extent MG1 and MG2 (Fig. 2a).

**Figure 2.**
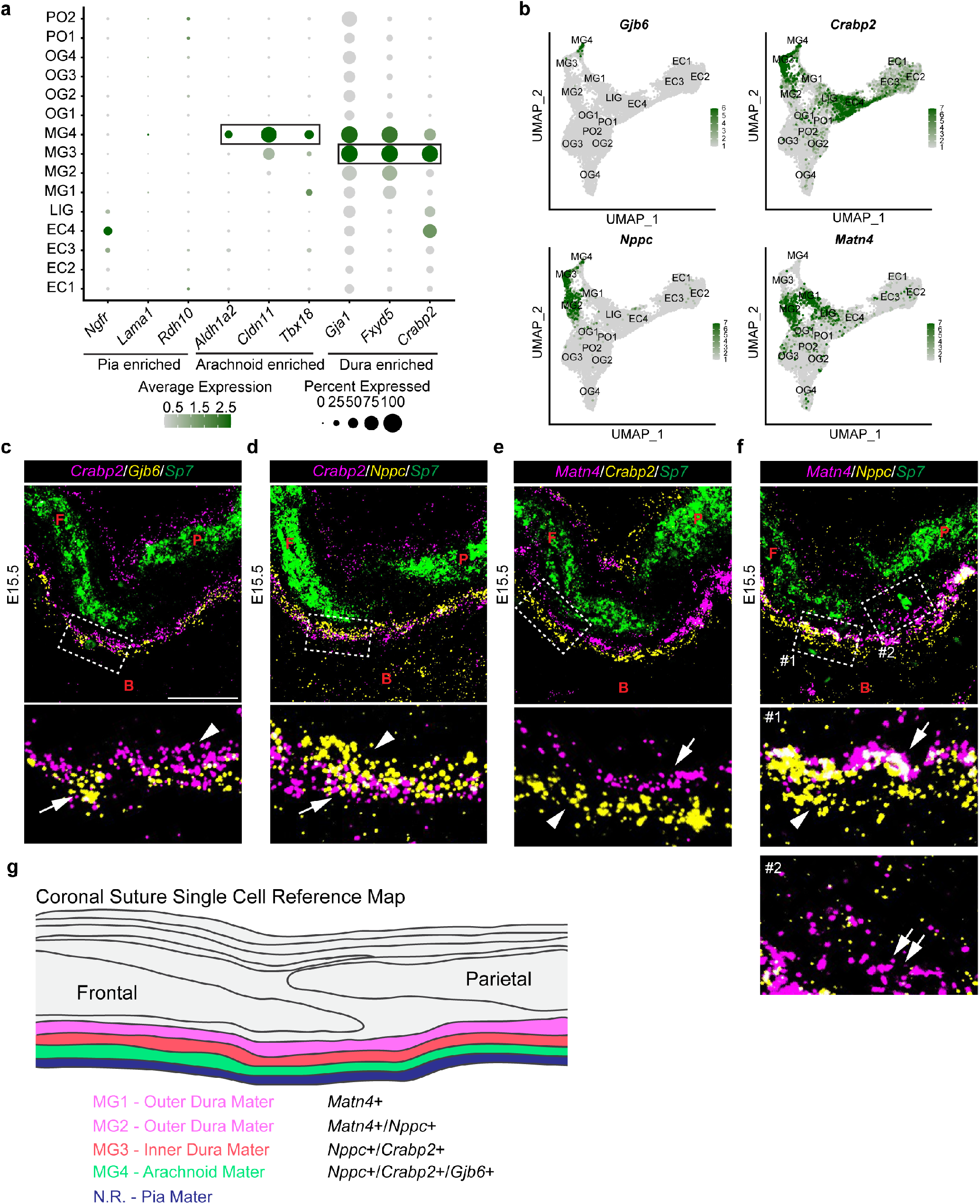
Diverse meningeal cell types are resolved by scRNA-seq. **a** Dot plot depicting selected markers previously associated with the pia mater, the arachnoid, and the dura mater. **b** Feature plots of genes validated by in situ experiments. **c-f** Combinatorial in situ analysis of coronal sutures for indicated markers at E15.5. *Sp7* marks the frontal (F) and parietal (P) bones, except for insets (boxed regions) below. **c** *Crabp2* and *Gjb6*. Arrow, Crabp2^+^/Gjb6^+^; arrowhead, Crabp2^+^. **d** *Crabp2* and *Nppc*. Arrow, Crabp2^+^/Nppc^+^; arrowhead, Nppc^+^. **e** *Matn4* and *Crabp2.* Arrow, *Matn4*^+^; arrowhead, *Crabp2*^+^. **f** *Matn4* and *Nppc*. Arrow, *Matn4^+^/Nppc^+^*; arrowhead, *Nppc*^+^; double arrows, *Matn4*^+^. **g** Model summarizing gene expression patterns of meningeal layers captured from single cell analysis. B, Brain. Scale bar = 50 μm.

To resolve the identity of MG4, we performed *in situ* experiments for a highly specific MG4 marker, *Gjb6*, in combination with the arachnoid/dura mater marker, *Crabp2* (Fig. 2b). *Crabp2* and *Gjb6* overlapped below the bone with *Crabp2* single-positive cells (MG3) found above the *Crabp2*^+^/*Gjb6*^+^ domain (Fig. 2c). Immunofluorescence for Crabp2 and Gja1 at E17.5 confirmed that these arachnoid/dura mater markers are excluded from the pia mater (Supplementary Fig. 3a). *Rgs5*+ pericytes were interspersed with *Gjb6+* cells in the MG4 arachnoid layer, consistent with the prominent vasculature extending from the border of the dura mater and through the arachnoid mater to the pia mater^26^ (Supplementary Fig. 3c). In the *Crapb2*^+^/*Gjb6*^−^ layer (MG3), we also observed co-expression of *Crabp2* with *Nppc*, with a zone of *Nppc*^+^/*Crabp2^−^* cells (MG2) above (Fig. 2d). In addition, *Nppc* was co-expressed with the known dura mater marker *Fxyd5^25^* (Supplementary Fig. 3b, d). MG3 may represent the dural border cell layer that has been described using EM in adult meninges^27,28^.

The *Nppc*^+^/*Crabp1*^−^ population above *Nppc*^+^/*Crabp2^+^* cells is consistent with MG2 identity in our UMAP analysis (Fig. 2b, d). This domain was also positive for *Matn4* and *Ctgf*, markers that only overlap with *Nppc* in MG2 (Fig. 2f, Supplementary Fig. 3b, e). *Matn4* expression does not overlap with the MG3/MG4 marker *Crabp2* (Fig. 2e), and chondrocytes in other sections express high levels of *Matn4* but not *Ctgf* (Supplementary Fig. 3f). We also detected some *Matn4* cells that were low or absent for *Nppc*, and these were situated closer to the suture and calvarial bones than the *Nppc*-high cells (Fig. 2f). Our UMAP analysis indicates that these *Matn4*^+^/*Nppc*^−^ cells represent MG1 (Fig. 2b). MG1 expressed higher levels of the chondrocyte markers *Col2a1* and *Acan* than MG2, but not at the level seen in bona fide chondrocytes (Supplementary Fig. 3g). These findings are consistent with MG1/MG2 representing a periosteal dura mater population^29^ primed to form cartilage, with the MG1 population having a stronger chondrogenic-like program. These data reveal a diversity of mesenchyme cell types within the meninges, with those closest to the suture sharing features with chondrogenic cells and likely functioning as a specialized connective tissue bridging the calvarial bones to the meninges (Fig. 2g).

### Diversity of ectocranial layers above the coronal suture

Ectocranial mesenchyme guides the migration of early osteogenic cells and helps establish suture boundaries^7,21,22^. Consistent with analysis of the mouse frontal suture^30^, we identified clusters enriched for markers of the dermis (*Ly6a*, *Dpt*; EC1-3) (Fig 1g, Fig. 3a). *Ly6a* is expressed in hypodermis of neonatal mouse skin^31^ and we observe co-expression of *Ly6a* with additional EC1-3 markers, *Pi16* and *Dpt*, within a single layer of mesenchyme above the skull bones (Fig. 3b, c; Supplementary Fig. 4a, b). Our previous work had identified expression of *Jag1* within both an ectocranial layer and suture mesenchyme cells^22^, and scRNAseq analysis shows high expression of *Jag1* in EC1, and to a lesser extent EC2 and EC3 (Fig. 1f-g, Supplementary Fig 4g). Co-expression of *Jag1* with *Pi16* confirms *Jag1* expression within the hypodermal layer (Supplementary Fig. 4h).

**Figure 3.**
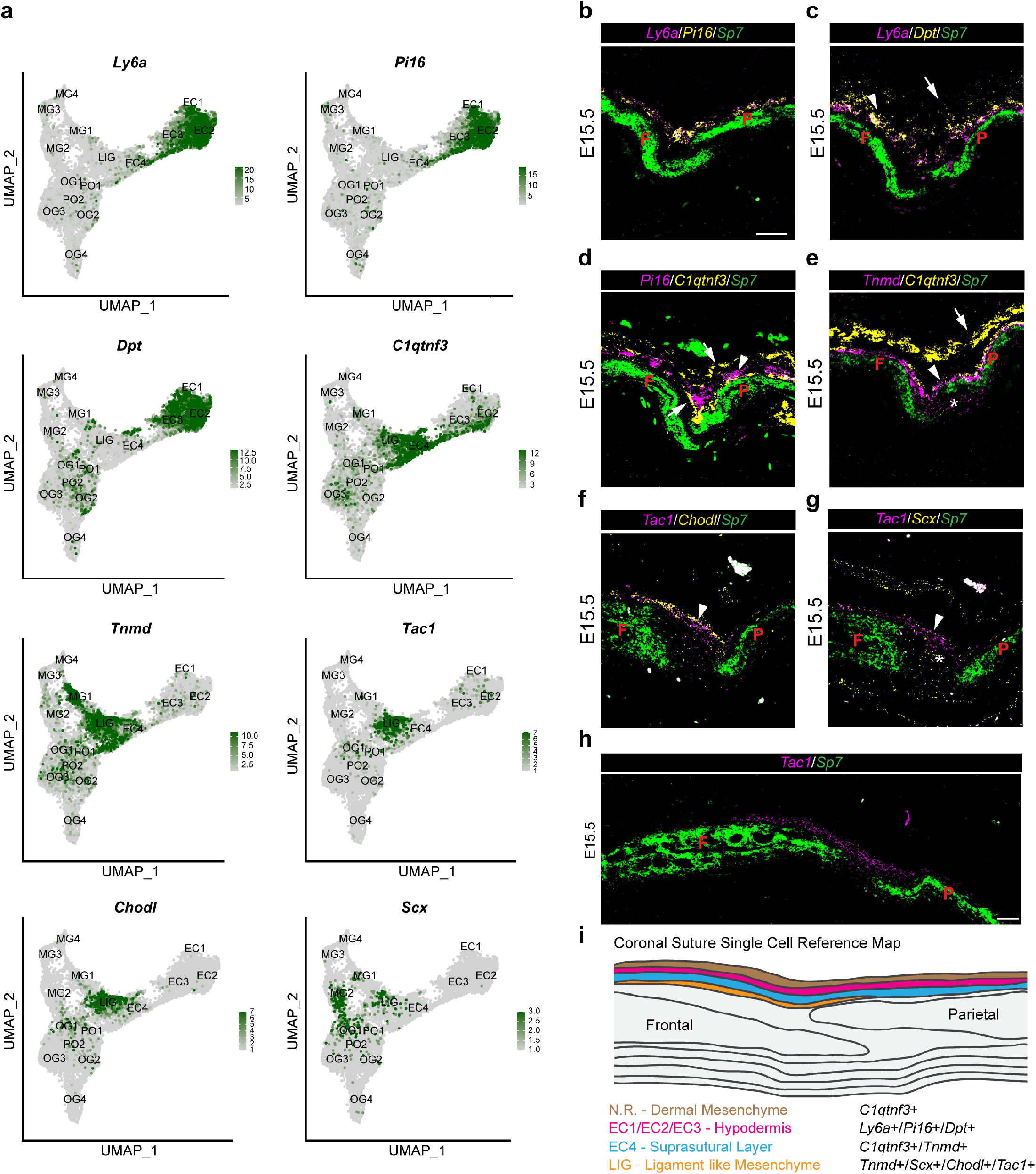
Multiple ectocranial layers overlay the coronal suture. **a** Feature plots of genes validated by in situ experiments. **b-h** Combinatorial in situ analysis of coronal sutures for indicated markers at E15.5. *Sp7* marks the frontal (F) and parietal (P) bones. **b** *Ly6a* and *Pi16*. **c** *Ly6a* and *Dpt*. Arrow, *Dpt*^+^; arrowhead, *Ly6a*^+^/*Dpt*^+^. **d** *Pi16* and *C1qtnf3.* Arrows, *C1qtnf3*^+^ layers; Arrowhead, *Pi16*^+^ layer. **e** *Tnmd* and *C1qtnf3.* Arrow, *C1qtnf3*^+^; arrowhead, *Tnmd*^+^/*C1qtnf3*^+^. Asterisk, *Tnmd* expression in MG1. **f** *Tac1* and *Chodl*. Arrowhead, *Tac1*^+^/*Chodl*^+^ ligament-like population. **g** *Tac1* and *Scx*. Arrowhead, *Tac1*^+^/*Scx*^+^ ligament-like population; asterisk, *Tac1*^+^*/Scx*^+^ suture mesenchyme. **h** *Tac1*. **i** Model summarizing gene expression patterns of ectocranial layers captured from single cell analysis. Scale bars = 50 μm.

In addition to hypodermal EC1-3 layers, we also noted a prominent EC4 population defined by *C1qtnf3* and *Tnmd* expression in our UMAP analysis (Fig. 3a). In situ experiments revealed two *C1qtnf3*^+^ ectocranial layers on either side of the *Pi16*^+^ hypodermis (Fig. 3d, Supplementary Fig. 4c). However, only the *C1qtnf3*^+^ layer between the calvarial bones and hypodermis expresses *Tnmd*, suggesting that this corresponds to EC4, which we term “suprasutural mesenchyme” (Fig. 3a, e; Supplementary Fig. 4d). It is likely that the outer *C1qtnf3*^+^ layer was removed along with the skin during dissection and thus not included in our single-cell analysis. Consistently, *Epha4*, which has also been shown to display ectocranial expression^7^, was most strongly expressed in this outer *C1qtnf3*^+^ layer despite not being particularly enriched in clusters EC1-4 in UMAP analysis (Supplementary Fig. 4i). Thus, *Jag1* and *Epha4* appear to label distinct ectocranial layers.

Interestingly, just underneath the *C1qtnf3*^+^/*Tnmd*^+^ layer (EC4), and sitting on top of the suture, we observed a *C1qtnf3*^−^/*Tnmd*^+^ population (LIG) that was enriched for a number of genes associated with tendon/ligament development (e.g. *Scx*, *Tnmd*, *Mkx, Thbs2, Bgn*), as well as markers of smooth muscle (e.g. *Acta2*, *Tagln*, *Myl9*) (Supplementary Fig. 4j). Two of the most specific markers for LIG were *Tac1,* a gene that encodes four neuropeptides including Substance P that is implicated in tendon mechanosensation^32^, and *Chodl*, a membrane protein with expression in tendons of humans^33^ and mouse limbs^34^ (Fig. 3a). In situ hybridization for *Tac1* and *Chodl* revealed highly restricted expression in mesenchyme overlying the coronal suture and connecting the edges of the frontal and parietal bones (Fig. 3f, h, Supplementary Fig. 4e). Although broader, in situ experiments showed that *Scx* and *Tnmd* were also expressed within this layer (Fig. 3g; Supplementary Fig. 4f, k), and we observed only minimal overlap between *C1qtnf3* and *Tac1* expression (Supplementary Fig. 4l). Thus, the LIG cluster represents a transcriptionally distinct and spatially confined subset of the ectocranium overlying the coronal suture and connecting the adjacent bones.

### Distinct osteoblast trajectories in the suture versus periosteum

The suture is a source of new osteoblasts for skull bone expansion. To better understand the trajectories of osteogenesis within and around the sutures, we identified six osteogenic clusters (OG1, OG2, OG3, OG4, PO1, PO2) based on expression of the osteoblast transcription factors *Runx2* and *Sp7*^35,36,37^, as well as *Dlx5 and Dlx6*^38,39,40^, (Supplementary Fig. 5). Pseudotime analysis of these clusters using Monocle3 revealed OG1 to be the earliest lineage cells, followed by two proliferative branches (PO1, PO2) and then distinct OG2 and OG3 trajectories that converged on the osteoblast cluster OG4 (Fig. 4a, b). These pseudotime trajectories were reflected by increased expression of *Runx2* and *Sp7* from OG1 to OG4 and accumulation of the mature osteoblast markers *Ifitm5* and *Dmp1* in OG4 (Fig. 4c). Prominent cluster markers included *Erg* and *Pthlh* (OG1), *Lef1* and *Inhba* (OG2)*, Mmp13* and *Podnl1* (OG3), and *Ifitm5*, *Dmp1*, and *Sost* (OG4) (Fig. 4e). Pseudotime visualization shows *Erg* (OG1) appearing earliest and shutting down as *Lef1* (OG2) and *Mmp13* (OG3) become expressed, followed by their extinguishment and progressive appearance of *Ifitm5, Dmp1,* and *Sost* (OG4) (Fig. 4d).

**Figure 4.**
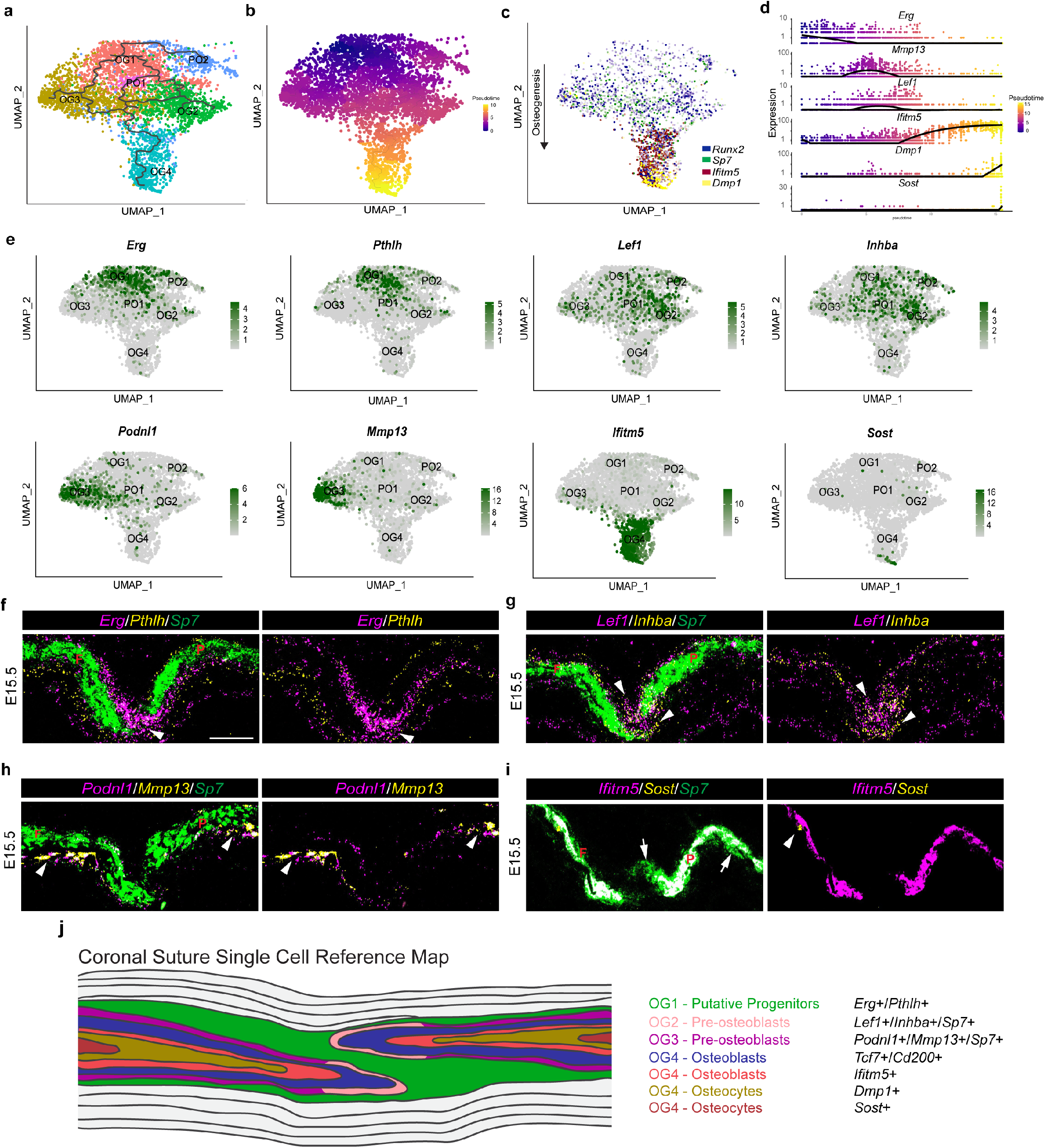
scRNA-seq captures various subtypes of osteoblasts at the coronal suture. **a** Lineage analysis and **b** Pseudotime analysis of osteoblast subset using Monocle 3. **c** Feature plot of selected markers of osteogenesis within the osteoblast subset. **d** Expression plots of selected genes across pseudotime. **e** Feature plots of genes validated by in situ experiments. **f-i** Combinatorial in situ analysis of coronal sutures for indicated markers at E15.5. *Sp7* marks the frontal (F) and parietal (P) bones in the left image of each pair. **f** *Erg* and *Pthlh.* Arrowhead marks suture mesenchyme. **g** *Lef1* and *Inhba*. Arrowheads mark *Lef1^+^/Inhba^+^* expression near bone tips. **h** *Podnl1* and *Mmp13.* Arrowheads mark *Podnl1^+^/Mmp13^+^* expression in periosteum distant from suture. **i** *Ifitm5* and *Sost*. Arrows, *Sp7^+^* expression in presumptive newly formed osteoblasts; arrowhead, *Sost*^+^. **j** Model summarizing gene expression patterns within the developing bones and coronal suture mesenchyme. Scale bar = 50 μm.

In situ validation at E15.5 showed co-expression of OG1 markers *Erg* and *Pthlh* in suture mesenchyme and extending along the edges of the frontal and parietal bone tips (Fig. 4f, Supplementary Fig. 6a). Interestingly, *Erg* but not *Pthlh* was asymmetrically distributed along the bones, with stronger expression above the frontal and below the parietal bone. *Gli1* and *Prrx1* showed similar asymmetric expression above the frontal bone, and *Gli1* and *Six2* below the parietal bone (Supplementary Fig. 7a-c). The different asymmetric mesenchyme patterns were not attributable to disproportionate populations of mesenchyme above and below the bones, as ectocranial and meningeal layers were generally evenly distributed along the bones (Figs. 2, 3). These findings highlight asymmetric distribution of the earliest osteogenic cells around the bone fronts, which may play a role in ensuring the later reproducible overlap of the parietal over the frontal bone.

We next examined expression of markers for OG2 and OG3, as pseudotime analysis suggested that these represent alternative pathways to osteoblasts (Fig. 4a). Markers for both OG2 (*Lef1*, *Inhba*) and OG3 (*Podnl1*, *Mmp13*) displayed expression along the outer domains of the *Sp7*^+^ bone surface, consistent with a pre-osteoblast identity. Whereas *Lef1*^+^/*Inhba*^+^ cells were most abundant at the edges of the growing bones near the suture, *Podnl1^+^*/*Mmp13*^+^ cells were enriched on bone surfaces further away from the suture (Fig. 4g, h; Supplementary Fig. 6b, c). OG2 may therefore represent more specialized suture-resident pre-osteoblasts, and OG3 periosteal pre-osteoblasts more generally found on bone surfaces.

Clusters PO1 and PO2 are enriched for markers of proliferation (*Mki67*, *Cenpf*, *Top2a;* Supplementary Fig. 6e), as well as expression of genes from suture-resident clusters, for example *Erg* (OG1) and *Lef1* and *Inhba* (OG2) (Fig. 1a, Fig. 4e). In situ validation revealed *Cenpf* expression around the tips of the growing bones and extending into the sutures, though in comparison to *Erg* it appeared largely absent from the central part of the suture (Supplementary Fig. 6f). These data suggest that proliferative pre-osteoblasts are concentrated at the bone tips, consistent with previous direct evidence for proliferative osteogenic cells at the leading edges of the calvarial bones^20^.

OG4 represents mature osteoblasts and osteocytes, as revealed by markers such as *Ifitm5*, *Dmp1* and *Sost* and its terminal position in the pseudotime analysis^41^ (Fig. 4a, c). We observe co-expression of *Sp7* with *Ifitm5* throughout most of the bone, except for the leading tips and the surfaces of bones which are *Sp7*-positive only, consistent with bone growth from both the leading edges and the periosteal surfaces (Fig. 4i, Supplementary Fig. 6d). Similarly, we observe protein expression of the osteoblast marker Cd200 in bone and the Wnt-responsive transcription factor Tcf7, a pre-osteoblast marker, along the bone tips and periosteal surfaces^43,44,45^, consistent with *Cd200* expression in cluster OG4 and *Tcf7* expression in both OG4 and the suture-specific pre-osteoblast cluster OG2 (Fig. 1g, Supplementary Fig. 8a-c). The mature osteocyte marker *Sost*, which labels the oldest cells in pseudotime, was expressed in only a few osteocytes distant from the suture (Fig. 4i). The spatial organization between osteoblasts, pre-osteoblasts, and progenitors was confirmed with double in situs between *Podnl1* and *Pthlh* or *Ifitm5* (Supplementary Fig. 6g-h). These findings are consistent with bone elongation occurring through osteogenesis at the suture and bone tips, with a distinct osteogenic pathway in the periosteum contributing to bone thickening (Fig. 4j).

### Signaling interactions between mesenchymal layers captured from single-cell analysis

To gain insights into potential signaling between osteogenic cells and adjacent ectocranial and meningeal mesenchyme, we surveyed our datasets for expression of ligands and receptors (Fig. 5a-b). In OG4 osteoblasts, we observed enrichment of *Tgfb1*, *Bmp3*, *Bmp4*, *Ihh*, and *Pdgfa*, with Ihh^46^ and Pdgfa^47,48^ known to be secreted by osteoblasts. In addition to osteoprogenitors (OG1), *Pthlh* ligand was expressed in several meningeal clusters. In the ectocranial clusters EC4 and LIG, which are situated closest to the suture, we detected enrichment of *Tgfb3*, *Fgf9*, *Fgf18*, *Wnt5a*, *Wnt9a*, *Wnt11*, and *Igf1* ligands. In the meningeal clusters closest to the suture (MG1, MG2), we also detected *Tgfb3* and *Igf1* ligand expression, as well as *Tgfb2* that has previously been shown to be a critical meningeal-derived factor for suture morphogenesis^49^. Reciprocally, we detected expression of the Tgfβ receptor *Tgfbr3*, the Fgf receptors *Fgfr1* and *Fgfr2*, the Wnt receptors *Fzd1*, *Fzd2*, and *Fzd6*, the Ihh *receptor Ptch1*, the Pthlh receptor *Pth1r*, and the Pdgfa receptor *Pdgfra* in various osteogenic clusters (Fig. 5b). Interestingly, *Fgfr1* was expressed more strongly in suture-resident pre-osteoblasts (OG2) and *Fgfr2* in periosteal pre-osteoblasts (OG3). To capture signalling interactions in an unbiased approach, we interrogated our data using the CellPhoneDB package^50^, scoring the ligand-receptor pairings between all clusters (Fig. 6c). Interactions between osteogenic clusters were notably weak, highlighting potential roles for osteogenic - non-osteogenic interactions in coronal suture regulation. Consistently, we noted that the mesenchymal populations closest to the suture (EC4 above, MG1/MG2 below) had the strongest interactions with osteogenic cells. The proliferative PO1 and pre-osteoblast OG3 clusters appeared to have the strongest connections with the surrounding mesenchyme, consistent with these populations residing at the surfaces of bones. Of the individual ligand-receptor interactions that were identified between EC4/MG1/M2 and osteogenic clusters, several pathways are relevant to coronal suture development, including Fgf, Tgfβ, and Wnt signalling^51^ (Supplementary Fig. 9).

**Figure 5.**
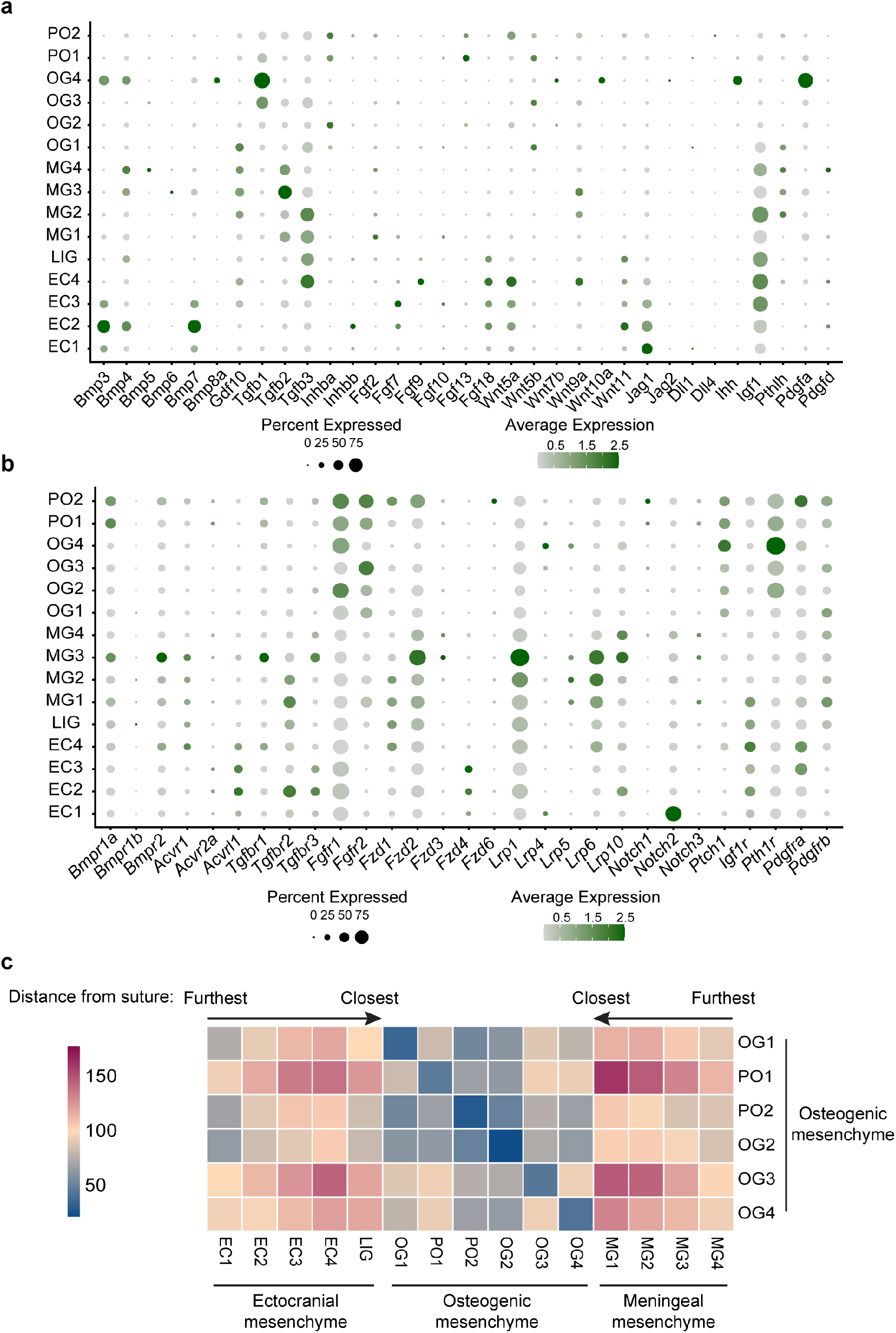
Ligand and receptor expression and predicted interactions at the coronal suture. **a** Dot plot of a selected group of secreted factors expressed within the coronal suture. **b** Dot plot of a selected group of receptors expressed within the coronal suture. **c** Heatmap of interaction scores between clusters from CellPhoneDB analysis. Ectocranial and meningeal clusters are arranged based on their validated distance from the coronal suture and bones. Heatmap scale shows counts of interactions.

**Figure 6.**
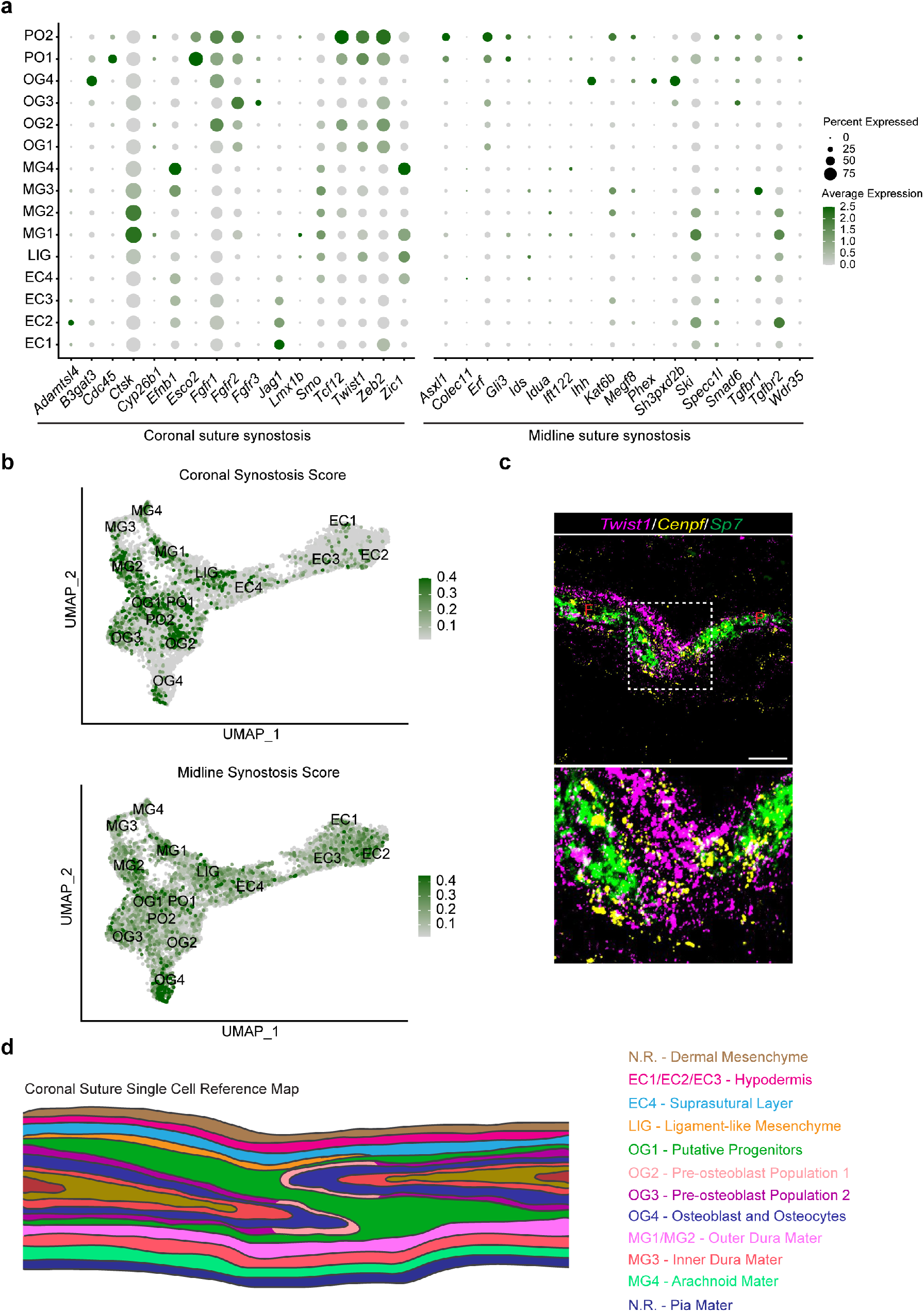
Genes associated with synostosis are expressed across multiple cell types in the coronal suture. **a** Dot plot of genes associated with coronal and midline suture synostosis. **b** Module score for coronal or midline synostosis genes plotted on to the osteogenic/mesenchymal subset UMAP. **c** Double in situ analysis of *Twist1* and *Cenpf* expression in the coronal suture relative to the frontal (F) and parietal (P) bones marked by *Sp7* expression (dashed box shows magnified region of suture mesenchyme below). **d** Model summarizing coronal suture cell types. Scale bars = 50 μm.

### Synostosis related genes display diverse expression patterns

To determine if genes underlying craniosynostosis in humans^51,52^ coalesce within a specific cell type, we mapped coronal and midline synostosis genes onto our dataset. Genes associated with midline suture synostosis were less abundant, suggesting that divergent gene expression patterns may contribute to suture-specific fusions (Fig. 6a). We noted that 8/17 coronal synostosis genes were selectively enriched in PO1 or PO2, suggesting misregulation of proliferating osteoblasts as a common mechanism of synostosis. By scoring every cell for the average expression of each coronal or midline synostosis gene, we noted that coronal synostosis scores were more restricted to the meningeal and osteogenic clusters, while midline synostosis genes were broadly distributed across clusters (Fig. 6b). Coronal synostosis genes with particular enrichment in PO1 and PO2 include *Fgfr1*, *Fgfr2*, *Tcf12*, *Twist1*, and *Zeb2*. Consistently, in situ validation revealed expression of *Twist1* in suture mesenchyme and cells lining the bone fronts, with *Twist1* expression also overlapping with the proliferative pre-osteoblast marker *Cenpf* (Fig. 6c, Supplementary Fig. 10). Selective enrichment of *Twist1* and *Tcf12* in proliferative pre-osteoblasts is consistent with functional studies showing roles for these transcription factors in regulating the balance between bone differentiation and proliferation^9,20^.

## Discussion

Proper formation of the coronal suture is a complex process requiring the coordinated development of multiple tissue types, which is reflected in the diverse genetic heterogeneity underlying coronal synostosis. Understanding the mechanisms that cause pathogenic suture fusion in distinct syndromic forms of synostosis has been limited by our incomplete knowledge of the cellular diversity of the developing coronal suture. In an effort to bridge this gap, we have generated a comprehensive spatial and transcriptomic map of the cell types that comprise and support the embryonic coronal suture (Fig. 6d).

The relationship of the embryonic progenitors that build the coronal suture with the adult sutural stem cells that grow the calvarium remains debated. Recent work on postnatal sutures has demonstrated that the suture mesenchyme houses resident skeletal stem cells that produce osteoblasts to grow skull bones^16,17,18,54^. However, these markers (*Gli1*, *Prrx1*, *Axin2*) broadly label mesenchyme throughout the embryonic skull, making them unsuitable for identifying osteogenic cells at earlier stages. Here we identify *Erg* and *Pthlh* as two of the earliest markers for the putative osteoblast progenitors concentrated in the embryonic coronal suture. As *Pthlh* is a marker for chondrocyte stem cells in the femoral growth plate^55^, Pthlh signaling may be a more general regulator of skeletal progenitors. As opposed to the adult skull where stem cells are tightly restricted to suture mesenchyme, embryonic *Erg^+^/Pthlh^+^* progenitors extend away from the suture along the surfaces of the frontal and parietal bone tips. Interestingly, these progenitors are asymmetrically distributed, with more cells along the lateral surface of the frontal bone and medial surface of the parietal bone. This asymmetric organization may ensure the reproducible architecture of the coronal suture, with the parietal bone consistently overlapping above the frontal bone. In a companion study, we have found that the asymmetric distribution of these progenitors, as marked by *Six2* and Grem1 protein, is lost upon disruption of *Tcf12* and/or *Twist1* function, consistent with bones meeting end-on-end and fusing in this craniosynostosis model (Ting et al. submitted).

Within the developing bone, we resolved a spatial hierarchy of osteoblasts, with progenitors located in the suture mesenchyme and giving rise to osteoblasts that are found progressively further away from the suture as they mature. Our data also revealed distinct routes of osteoblast differentiation through either suture-resident pre-osteoblasts or periosteal cells more broadly distributed along the surfaces of bones. Suture-resident pre-osteoblasts are enriched for Wnt-related genes, perhaps reflecting the known involvement of genes from this pathway in coronal suture development^56,17^, as well as proliferative markers such as *Cenpf*. These findings support a model in which proliferative bone growth from the coronal suture extends the lengths of bones, whereas more restrained osteoblast differentiation along the bony surfaces increases the thickness of the calvarial bones, particularly at postnatal stages^57^.

The underlying meninges are important in sutural and calvarial development^12,58^, and we uncovered multiple distinct layers associated with the coronal suture. In particular, we resolved the dura mater into a dural border cell layer and two layers consistent with periosteal dura^27,28^. Interestingly, the periosteal dura layer, which sits closest to the suture and calvarial bones, was distinguished by expression of several chondrogenic markers, including *Matn4*, *Ctgf*, and lower levels of *Col2a1* and *Acan*. Cartilage formation beneath sutures has been linked to normal and pathological suture fusion^59,60^, yet the precise source of such cartilage has remained unclear. It will be interesting to assess whether the periosteal dura layers contribute to the natural and ectopic cartilage formation associated with suture closure. In addition, the periosteal dura layers express multiple signalling factors implicated in suture regulation, including *Tgfb2*, *Fgf2*, *Gdf10,* and *Ctgf*^61^, and CellPhoneDB analysis points to signalling interactions between the periosteal dura layers and pre-osteoblasts. Our single-cell atlas therefore provides testable hypotheses for how specific meningeal cell types may regulate calvarial bone formation in a paracrine fashion.

The ectocranial mesenchyme above the coronal suture is also known to regulate suture patency^7,22^, and here we identify at least four distinct layers. The two outermost layers likely represent the lower limits of the skin, including dermal reticular fibroblasts (*C1qtnf3*^+^) and non-osteogenic *Ly6a^+^* (*Sca1*^+^) positive hypodermis^31,62,63^. Whereas the hypodermis expresses *Jag1*, *Epha4* is enriched in the dermal reticular fibroblast layer. As both *Jag1* and *Epha4* have been implicated in coronal suture formation^7,22^, our data suggest that multiple ectocranial layers play a role in suture regulation. Below the hypodermis, we identified a *C1qtnf3*^+^/*Tnmd*^+^ layer that we termed suprasutural, as well as a related and tightly associated layer enriched for expression of genes involved in ligament formation, cellular contraction, and mechanosensation. A dense network of collagen fibres is present within adult mouse coronal sutures^64^ and may correspond to the ligament-like population in our embryonic dataset. One possibility is that this ligament-like population, which connects the lateral tips of the frontal and parietal bones, may contribute to the flexibility of the sutures, for example to accommodate compression of the calvarium during birth. More speculatively, this population could also function to interpret mechanical forces and transmit these to the osteogenic cells within the suture, thus coupling expansion of the brain to calvarial growth. As with the periosteal dura layers of the meninges, the suprasutural and ligament-like layers have strong predicted signalling interactions with osteogenic cells within the suture, including expression of members of the Tgfβ, Fgf, Wnt, and Igf1 signalling families. It will be interesting to determine the extent to which meningeal and ectocranial layers regulate osteogenic differentiation through similar or distinct means, and whether these layers help sense mechanical forces driving skull expansion.

The broad distribution of craniosynostosis-related genes within our dataset suggests diverse etiologies of synostosis. However, we did note that nearly half of synostosis genes had highest expression within the proliferative osteogenic clusters, including *Twist1* and *Tcf12* that have been implicated in negative regulation of the rate of proliferative bone growth in the calvarium^20^. Thus, misregulation of osteogenic cell proliferation may be a common driver of coronal synostosis, although other coronal synostosis genes had preferential enrichment in meningeal (*Ctsk*, *Efnb1*) or ectocranial (*Jag1*) layers. In addition, midline synostosis-related genes had much lower expression within our dataset, which may reflect the distinct genetic sensitivities of particular cranial sutures.

Comparison to a recently published dataset for the frontal suture^30^ highlights both conserved and divergent features of the coronal suture (Supplementary Fig. 11). Similar hypodermis (EC1-3 <-> HD), dura mater (MG1-4 <-> DM), and osteoblast (OG4 <-> OB) populations were identified within both. A subpopulation of *Ctgf*-expressing FS3 cells was also found below the frontal suture, which may correspond to MG1 in our coronal suture dataset. Similar populations to EC4 (FS1) and LIG (FS2) were also captured within the metopic suture datasets. However, there are prominent differences in the spatial distribution of these clusters between sutures. Whereas the EC4 suprasutural population is distributed above the bones in the coronal suture, the comparable FS1 population appears to occupy the bulk of the suture mesenchyme in the frontal suture. Similarly, the LIG population in the coronal suture connects adjacent bones, as true ligaments do, yet the comparable FS2 population is embedded within the frontal suture mesenchyme distant from bone. The biggest differences between the frontal and coronal datasets can be seen in the osteogenic mesenchyme. Whereas both datasets capture osteoblasts and proliferating pre-osteoblasts (PO1/PO2 <-> FS4), the earliest *Erg*^+^ progenitors and distinct routes through suture-resident pre-osteoblasts (OG2) and periosteal pre-osteoblasts (OG3) at the coronal suture were not apparent in the frontal suture dataset. The frontal suture cluster FS3 appears closest to these signatures, and it is possible that periosteal tissues were not included to the same extent in dissection of the frontal suture. However, there is no evidence of the asymmetric distribution of osteoprogenitors seen at the coronal suture for the frontal suture, which likely reflects that bones meeting at the frontal suture do not overlap in the same way as at the coronal suture. Our analysis therefore reveals differences in the topological arrangements of cell populations between sutures, and possibly in the identities and functions of the populations themselves. This atlas of the coronal suture therefore will be an important resource for understanding why specific sutures are reproducibly affected in distinct human craniosynostosis syndromes.

## Methods

### Coronal Suture Dissociations

E15.5 and E17.5 embryos were isolated and the bony skull was dissected away from the skin and brain. For E15.5 embryos the coronal suture was dissected away from the skull cap using microdissection scissors, and 10 pooled coronal sutures were rinsed in PBS and enzymatically dissociated with a final concentration of 3 mg/mL of Collagenase II (Worthington) and 4 units/mL of Dispase (Corning) in DMEM/F12 (Corning) for 45 min. For E17.5 embryos coronal sutures were dissected out in ice cold PBS using a scalpel blade, isolating a strip containing the overlapping frontal and parietal bone fronts (which appears opaque compared to adjacent regions) and avoiding the most apical and basal aspects of the suture. Isolated sutural strips from embryos from two litters in 3 batches (batch 1, 10 sutures from litter 1; batch 2 and 3, 3 sutures each from litter 2) were cut into small fragments in HBBS and digested using Collagenase IV (Worthington, USA; final concentration in HBBS of 2 mg/mL) for 30 min. Dissociation was terminated with 2% Fetal Bovine Serum and cells were passed through a 0.35 μM filter (E15.5) or Pluri-strainer Mini 70 μm (E17.5; pluriSelect Life Science, Germany). For E15.5 sample preparation, dead cells were removed using the Dead Cell Removal kit (Miltenyi Biotec 130-090-101) and cells counts were determined with a hemocytometer. The three batches of E17.5 dissociated cells were separately sorted by FACs to remove debris, cell doublets and likely dead cells (BD FACSAria Fusion; 100 μM nozzle) prior to library preparation.

### scRNA-seq library preparation and sequencing

Transcriptome libraries for single cells were captured using 10X genomics Chromium Single Cell 3’ Library and Gel Bead Kit v2 following manufacturer’s guidelines. Sequencing for E15.5 coronal sutures was performed with Illumina’s HiSeq 3000/4000 PE Cluster Kit at the Children’s Hospital Los Angeles’ Molecular Genomics Core, and for E17.5 cells, PE sequencing was run on an Illumina HiSeq 4000 at the Oxford Genomics Centre Wellcome Centre for Human Genetics, Oxford achieving an average of ~150,000 mean reads per cell for E15.5 and ~92,000 mean reads per cell for E17.5.

### Bioinformatics analysis

Quality control of raw reads was performed with FastQC version 0.11.7, fastq_screen version 0.7.0 and multiqc^65^ version 0.9. Samples were counted individually and aggregated with cellranger (10X Genomics) version 2.1.1 using the mm10 mouse transcriptome. All other parameters were set to their default values. Data analysis was performed with Seurat^23^ version 3.2.0. The aggregate gene/barcode matrix (‘raw_gene_bc_matrices_mex’) was loaded using ‘CreateSeuratObject’ and ‘Read10X’ with ‘min.cells = 10, min.genes = 200’. Cells were filtered to exclude cells with fewer than 1000 UMIs, fewer than 1000 genes, more than 7.5% mitochondrial content (calculated as the fraction of reads assigned to a gene on the chromosome ‘MT’), and more than 3% Hbb/Hba centent (to remove red blood cells, and cells with high contamination for red blood cell specific genes). Filtered cells were normalized with SCTransform, and cell cycle scoring and regression was performed for each dataset with the default list of human cell-cycle genes (converted to mouse gene symbols). Datasets were integrated based on the tutorial “Integration and Label Transfer” from Seurat. In brief, an object list including both datasets was created, 3000 features were selected using the SelectIntegrationFeatures following by PrepSCTIntegration, anchors were identified using FindIntegrationAnchors, and the data was integrated using IntegrateData. Dimensionality reduction was performed using the first 30 principal components (PCs) with UMAP^66^ (‘min_dist = 0.5, n_neighbors = 50’). Clustering was performed with ‘FindClusters’ and ‘resolution=1’. Cluster marker genes were identified with ‘FindAllMarkers’ using parameters ‘min.pct = 0.2, logfc.threshold = 0.5, max.cells.per.ident = 1000, min.cells.gene = 5’. Secondary dimensionality reduction and clustering of the osteogenic and mesenchymal subset was performed as above, after subsetting for cluster numbers 0, 1, 2, 4, 5, 7, and 9. Trajectory analysis was carried out on the osteogenic subset (clusters OG1-4, PO1-2) using Monocle 3^67^. The Seurat integrated object was converted into a Monocle cell dataset and the cluster information and UMAP coordinates carried over from the Seurat object prior to following the Monocle3 recommended protocols. Cell–cell communication analysis was performed using CellphoneDB v2.1.4^50^, after transforming mouse genes to human homologs. We prioritised potential interactions (p-value < 0.01) and manually selected those that were of biological relevance. Synostosis scores was determined using AddModuleScore in Seurat.

### RNAScope and immunohistochemistry

RNAscope in situ hybridization was performed the RNAscope Multiplex Fluorescent Kit v2 (Advanced Cell Diagnostics, Newark, CA) according to the manufacturer’s protocol for fixed-frozen sections, with one modification. To retain optimal sectioning quality, the heat antigen retrieval was omitted. TSA® Plus (Fluorescein, Cy3 and. Cy5) reagents were used at 1:1000. For immunohistochemistry, wild-type C57BL/6 embryos were collected at E17.5 and rinsed in ice-cold PBS for 30 minutes followed by head dissection, skin removal and overnight incubation in 4% PFA at room temperature. Heads were then washed multiple times in PBS, decalcified for 2 hours in Calci-ClearTM Rapid (HS-105, National Diagnostics), dehydrated, and paraffin embedded. Embedded tissue was sectioned (5 μm) and then rehydrated, stained and visualized using ImmPRESS® HRP Anti-Rabbit IgG (Peroxidase) Polymer Detection Kit (MP-7451, Vector Laboratories). Primary antibodies were diluted in cold TBS and incubated overnight at 4°C. In order to detect primary antibodies other than those raised in rabbit, donkey anti-sheep/goat/rat IgG-HRP was used (all 1:200, A16041, sc-2020, A18739, respectively). The tissue sections were subsequently imaged using an Olympus BX60 Microscope (Olympus) and/or NanoZoomer 2.0 HT (Hamamatsu). Secondary antibodies include donkey anti-sheep IgG-HRP (A16041), donkey anti-goat IgG-HRP (sc-2020), and donkey anti-rat IgG-HRP (A18739). To study Tcf7/Cd200 and Tcf7/Dmp1 localization, double immunofluorescence staining was performed on 5 μm E17.5 sections. Following deparaffinisation, rehydration, and heat-mediated antigen retrieval in 10 mM sodium citrate buffer solution (pH 6), samples were blocked in 4% Donkey Serum (D9663, Sigma-Aldrich) for 30 min. Individual sections were then incubated overnight at 4 °C with a mixture of Tcf7 (1:200; C63D9, Cell Signaling Technology) and Cd200 (1:200; AF2724, R&D systems) or Tcf7 (1:200; C63D9, Cell Signaling Technology) and Dmp1 (1:400; AF4386, R&D systems) primary antibodies. Antigen detection was performed using appropriate combination of the Alexa Fluor 488, 555 and 647 secondary antibodies (all 1:500; A21206, A21432 or A11015, A31573; Thermo Fisher Scientific) for 2 h at room temperature in the dark. All primary/secondary antibodies were diluted in SignalBoost™ Immunoreaction Enhancer Kit (407207–1KIT, Calbiochem). After three washes in PBS, sections were incubated with DAPI (1 μg/mL) (Roche, cat # 10 236 276 001). Following multiple washes in PBS, slides were mounted using Vectashield® Antifade Mounting Medium (H-1000Vector Laboratories, Inc.). Imaging was performed using a Zeiss LSM 780 Upright Multi-Photon Confocal Microscope with LD LCI PA 25× /0.8 DIC WD = 0.57 mm Imm Corr (UV)VIS-IR (Oil-Immersion) and Plan-Apochromat 63x/1.4 Oil objectives. Images were obtained using ZEISS ZEN Microscope software. When using two rabbit antibodies, for example the co-localisation of Gja1 (1:100; 3512, Cell Signalling Technology) and Crabp2 (1:750; 10225-1-AP, Proteintech), we used the TSA Cyanine 3 Plus Evaluation Kit (NEL744E001KT, Perkin Elmer) following the manufacturers’ instructions. Primary antibodies were visualised using ImmPRESS® HRP Anti-Rabbit IgG (MP-7451, Vector Laboratories) followed by incubation with Fluorescein (FP1168) or Cyanine 3 (FP1170) Amplification Reagents (both 1:200) and imaged by a Zeiss LSM 780 Upright Multi-Photon Confocal Microscope (same parameters as above).

### Data availability

The RNAseq data are available in the GEO repository, accession: GSE163693.

## Acknowledgments

We thank Kevin Clarke and Craig Waugh from the WIMM Flow Cytometry Facility, and Claire Arata, Maxwell Serowoky, and Francesca Mariani from the University of Southern California for help with single-cell dissociation and isolation. We thank Julie Siegenthaler from the University of Colorado Anschutz Medical Campus for helpful advice and feedback. We thank the Oxford Genomics Centre at the Wellcome Centre for Human Genetics and the Children’s Hospital Los Angeles’ Molecular Pathology Genomics Core for next-generation sequencing. Work was supported by Wellcome (102731 to AOMW), Action Medical Research (GN2483 to SRFT), VTCT Foundation Fellowship (SRFT, AOMW), the MRC through the WIMM Strategic Alliance (G0902418 and MC_UU_12025), Burroughs Wellcome Trust (DTF), HHMI Hanna H. Gray Fellows Porgram (DTF), National Institutes of Health (R01DE026339 to JGC and REM). The views expressed in this publication are those of the authors and not necessarily those of funding sources.

## Contributions

D.T.F., A.O.M.W., R.E.M., S.R.F.T. and J.G.C. conceived and designed the study. D.T.F., Y.Z. N.K., N.A. and S.R.F.T. carried out the single-cell and bioinformatic analysis. D.T.F. performed the RNAScope validation experiments, and H.M. the immunolocalizations. S.R.F.T., A.O.M.W. and J.G.C. supervised the research. D.T.F., S.R.F.T. and J.G.C wrote the paper with contributions from all authors.

